# Coxsackievirus group B3 regulates ASS1-mediated metabolic reprogramming and promotes macrophage inflammatory polarization in viral myocarditis

**DOI:** 10.1101/2024.05.08.593129

**Authors:** Qiong Liu, Yinpan Shang, Ziwei Tao, Xuan Li, Lu Shen, Hanchi Zhang, Zhili Liu, Zhirong Rao, Xiaomin Yu, Yanli Cao, Lingbing Zeng, Xiaotian Huang

## Abstract

Coxsackievirus group B3 (CVB3) belongs to the genus *Enteroviruses* of the family *Picornaviridae* and is the main pathogen underlying viral myocarditis (VMC). No specific therapeutic is available for this condition. Argininosuccinate synthase 1 (ASS1) is a key enzyme in the urea cycle that converts citrulline and aspartic acid to argininosuccinate. Here, we found that CVB3 and its capsid protein VP2 inhibit the autophagic degradation of ASS1 and that CVB3 consumes citrulline to upregulate ASS1, triggers urea cycle metabolic reprogramming, then activates macrophages to develop pro-inflammatory polarization, thereby promoting the occurrence and development of VMC. Conversely, citrulline supplementation to prevent depletion can downregulate ASS1, rescue macrophage polarization, and alleviate the pathogenicity of VMC. These findings provide a new perspective on the occurrence and development of VMC, revealing ASS1 as a potential new target for the treatment of this disease.

**IMPORTANCE:** Viral myocarditis (VMC) is a common and potentially life-threatening myocardial inflammatory disease, most commonly caused by CVB3 infection. So far, the pathogenesis of VMC caused by CVB3 is mainly focused on two aspects: one is the direct myocardial injury caused by a large number of viral replication in the early stage of infection, and the other is the local immune cell infiltration and inflammatory damage of the myocardium in the adaptive immune response stage. There are few studies on the early innate immunity of CVB3 infection in myocardial tissue, but the appearance of macrophages in the early stage of CVB3 infection suggests that they can play a regulatory role as early innate immune response cells in myocardial tissue. Here, we discovered a possible new mechanism of VMC caused by CVB3, revealed new drug targets for anti-CVB3 and discovered the therapeutic potential of citrulline for VMC.

## INTRODUCTION

Myocarditis is an inflammatory disease of the myocardium caused by infections and autoimmune diseases. Viral infections are the most commonly reported causes of myocarditis, and the causative viruses tend to enter the myocardium^[1]^. Epidemiological studies have linked myocarditis to enteroviruses, especially Coxsackievirus group B3 (CVB3)^[2]^, and a United Kingdom statistic shows that 25% of viral myocarditis (VMC) cases are caused by CVB3 infection^[3]^. However, the mechanism by which CVB3 infection causes VMC is not well understood. Previous studies have suggested that after CVB3 infection, cardiac-resident cells, such as cardiomyocytes, release large amounts of pro-inflammatory cytokines and chemokines to establish a pro-inflammatory environment^[4]^. Thereafter, various innate immune cells and adaptive immune cells infiltrate tissue and promote tissue damage by secreting cytokines or via lysis functions^[5, 6]^. Ultimately, myocardial repair and fibrosis proliferation occurs^[7]^. Among the various immune cells, macrophages have been proven to play an important role in both early innate and adaptive immunity. First, normal myocardial tissue contains a small number of macrophages called cardiac-resident macrophages, which can perform immune surveillance functions in the first place^[8, 9]^. Second, macrophages are the main infiltrating cell subset in the early stages of VMC, promoting inflammatory damage^[10]^. Moreover, histological or immunohistological evidence of inflammatory cell infiltration is the gold standard for diagnosing myocarditis, and in VMC models, a significant increase in the number of macrophages in the myocardium is observed, which plays an important role in the progression of myocarditis^[11–13]^.

Macrophages are typical innate immune cells that play a key role in the body’s antiviral immune response. Macrophage polarization is not fixed; it is a plastic, dynamic, and tightly controlled process, regulated by various signals, and macrophages can often be divided into pro-inflammatory (M1-like) and anti-inflammatory (M2-like) macrophages according to their function^[14]^. The former mainly plays a role in the acute stage of the disease, promoting the occurrence and progression of inflammation and facilitating pathogen clearance but aggravating immune injury. The latter mainly plays a role in the disease recovery process and can inhibit inflammatory responses and promote tissue repair but is not conducive to pathogen clearance^[14]^. In CVB3-induced VMC, macrophages undergo pro-inflammatory polarization, which strongly induces the release of cytokines (interleukin-1, IL-1; interleukin-6, IL-6; Tumor necrosis factor alpha, TNF-α) and inducible nitric oxide synthase (iNOS), resulting in intense inflammation and increased myocardial damage^[15–17]^. The pro-inflammatory polarization of macrophages is essential for the activation of inflammation, and these cells are often studied as effector-inflammatory infiltrating cells of adaptive immunity after viral infection. However, their role as early innate immune response cells in myocardial tissue is little studied, and further research is needed to determine how macrophages recognize CVB3 to activate pro-inflammatory polarization in the innate immune stage.

Recently, many studies found that cellular metabolism may play an important role in macrophage polarization. Glucose, amino acids, and lipids all play key roles in macrophage polarization^[18–20]^. The urea cycle is a key process in the metabolism of amino acids in cells, mainly in the liver, where it is used to eliminate excess toxic ammonia from the body, and argininosuccinate synthase 1 (ASS1) is a key enzyme in the urea cycle that converts citrulline and aspartic acid to argininosuccinate^[21]^. In recent years, with the application of metabolomics, an increasing number of studies have demonstrated a correlation between the urea cycle and viruses, reporting that their interaction affects the disease process. In a mouse model of lymphocytic choriomeningitis virus (LCMV) infection, type I interferon signaling alters the downregulation of ornithine transcarbamylase (OTC) and ASS1, key enzymes of the urea cycle in hepatocytes, leading to systemic metabolic alterations, attenuating antiviral T cell responses, and improving liver injury^[22]^. Mao et al. found that the upregulation of ASS1 triggers citrulline depletion, leading to pro-inflammatory polarization in macrophages^[19]^. These studies suggest that ASS1 may also be involved in CVB3-induced pro-inflammatory macrophage polarization, which triggers urea cycle reprogramming. However, the mechanism of action of urea cycle reprogramming in the host response induced by CVB3 is unclear and deserves further exploration.

CVB3 belongs to the genus *Enteroviruses* of the family *Picornaviridae* and has a single-stranded positive-stranded RNA genome with a length of approximately 7396 bp^[23]^. Its genome encodes four capsid proteins, namely VP1, VP2, VP3, and VP4^[24]^. Among them, VP2 is an important pathogenic factor causing cardiomyocyte injury^[25]^. Some studies have shown that the viral capsid protein has a regulatory function in addition to maintaining the structure of the virus; however, the regulatory function of the viral capsid protein VP2 is currently unclear. Here, we focused on VP2, exploring its regulatory role regarding ASS1. We found that CVB3 and its capsid protein VP2 inhibit the fusion of autophagosomes with lysosomes, thus inhibiting the autophagic degradation of ASS1. In this study, we constructed a cellular model of CVB3-induced macrophage polarization and elucidated the relationship between CVB3 and amino acid metabolism reprogramming for the first time. We found an interaction between CVB3 capsid protein VP2 and ASS1 to upregulate ASS1, resulting in amino acid metabolism reprogramming, inducing macrophage pro-inflammatory polarization, and promoting the occurrence of VMC. These findings suggest that ASS1 may be a potential target for VMC treatment. We also observed the alleviating and therapeutic effects of the ASS1 substrate citrulline against VMC. Overall, this project provides a new research direction to understand the occurrence and development mechanisms of VMC.

## RESULTS

### CVB3 induces pro-inflammatory macrophage activation that promotes the development of VMC

To systematically investigate the mechanism by which CVB3 causes VMC, we established a CVB3-induced acute VMC model to study CVB3 infection. BALB/c mice (4 to 6 weeks old) were infected intraperitoneally with 5×10^7^ PFU of CVB3. The mice were euthanized and their hearts were collected and analyzed by immunofluorescence and hematoxylin-eosin (H&E) staining several hours post-infection. Although we found no obvious inflammation in the hearts of CVB3-infected mice, significant macrophage infiltration was observed, and the increase in macrophages was most pronounced in the early stages of infection, suggesting that macrophages may be involved in the early innate immune response to VMC (**Fig. 1A and Supplementary Fig. 1**). Currently, while macrophages are often studied in animal models of VMC, there is no suitable cell model to investigate molecular mechanisms^[26, 27]^. Therefore, we sought to explore the mechanisms of macrophage activation in depth to understand the molecular basis of VMC^[28]^.

**Figure 1.**
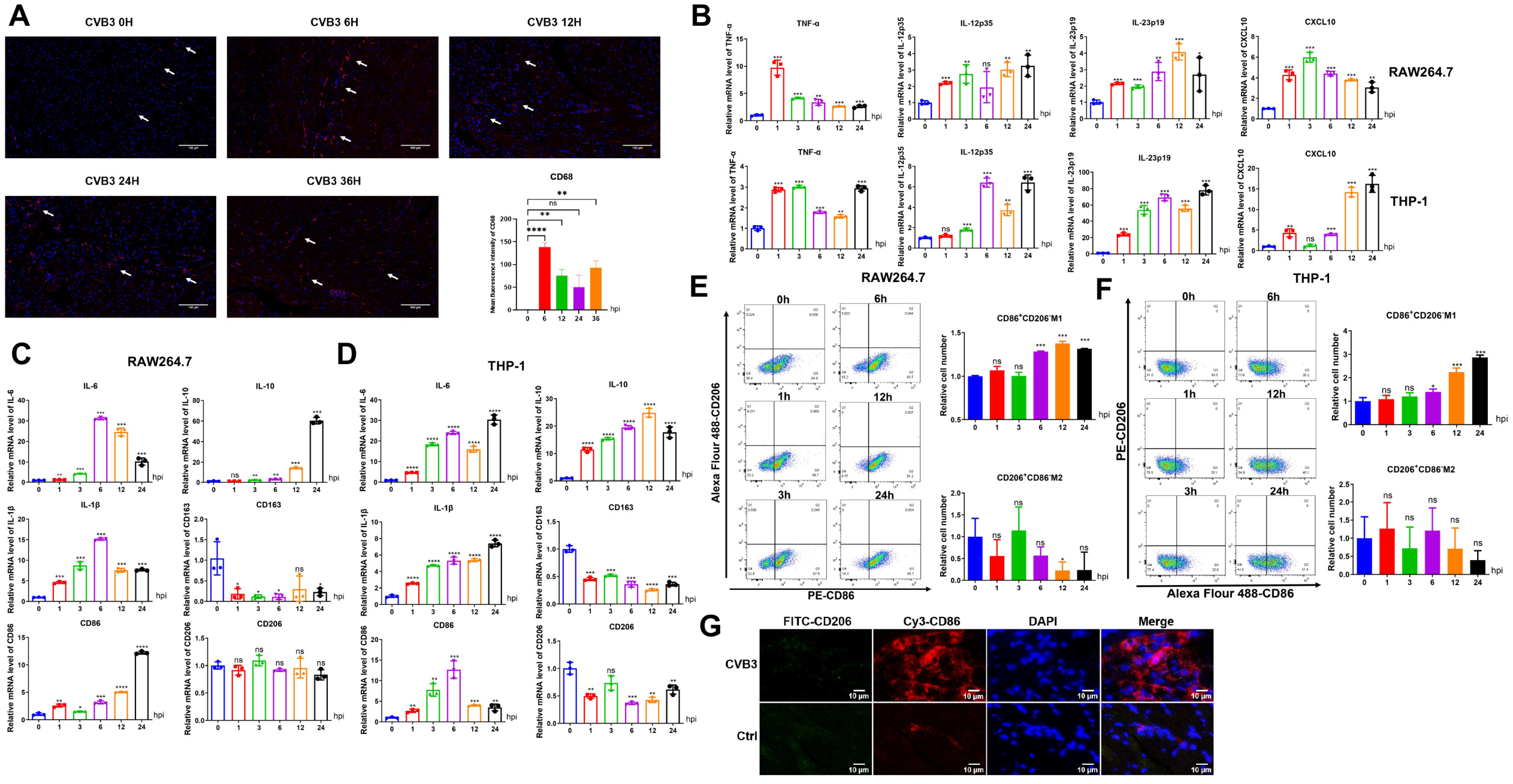
CVB3 induces pro-inflammatory macrophage activation. (A) Representative immunofluorescence staining of CD68 in heart sections of mice infected with 5×10^7^ PFU of CVB3 at several hours post-infection (hpi). Nuclei were stained with DAPI. Quantitative CD68 expression is presented as mean fluorescence intensity. Scale bar = 100 μm. n = 6 in each group. (B, C, and D) Expression levels of pro-inflammatory cytokine and M1/M2 macrophage marker mRNA after CVB3 (MOI = 20) treatment of macrophages, measured by RT-qPCR. (E and F) Flow cytometry was used to detect the changes of CD86^+^CD206^-^M1 macrophages incubated with CVB3 (MOI = 20) for different durations. (G) Immunofluorescence was used to detect the types of macrophage infiltration in mouse heart tissue with VMC. The expression of FITC-CD206 and Cy3-CD86 in mouse VMC was observed by fluorescence microscopy. The FITC-labeled proteins appear green, the Cy3-labeled proteins appear red, and nuclei appear blue. Values are shown as the mean ± SEM of three independent experiments. (*, P < 0.05; **, P < 0.01; ***, P < 0.001; ****, P < 0.0001; ns, P > 0.05).

To determine the appropriate experimental conditions for CVB3 to act on macrophages, RAW264.7 cells were infected or incubated with CVB3 (multiplicity of infection [MOI]=20). The results showed that the expression levels of VP1 completely differed between two methods. Over time, the expression of VP1 mRNA in CVB3-incubated RAW264.7 cells remained high (**Supplementary Fig. 2**). Monocyte-macrophage infiltration often occurs when a large number of cardiomyocytes have been broken. At this point, a large number of CVB3 virions replicating in cardiomyocytes are released, allowing the heart-resident macrophages to fully engage with the virus^[29]^. We speculated that the incubation treatment allowed macrophages to be continuously stimulated by CVB3, better simulating the real external environment of myocardial macrophages in CVB3-infected patients. Next, we investigated the role of CVB3 in macrophage activation. The expression levels of macrophage polarization markers (IL-6, IL-1β, IL-10, CD86, CD163, CD206) were measured by quantitative reverse transcription polymerase chain reaction (RT-qPCR) and flow cytometry. First, after CVB3 was incubated with macrophages, the cells showed higher levels of the pro-inflammatory cytokines and chemokines TNF-α, interleukin-12 p35 (IL-12 p35), interleukin-23 p19 (IL-23 p19), and C-X-C motif chemokine ligand (CXCL) 10 (**Fig. 1B**). In the meantime, RAW264.7 and THP-1 cells displayed a dramatic increase in cellular pro-inflammatory macrophage gene levels upon CVB3 incubation at various time points post-incubation, whereas anti-inflammatory macrophage gene levels remained unchanged (**Fig. 1C and D**). Flow cytometry data further confirmed these observations. The proportion of pro-inflammatory macrophages increased by varying degrees, while the proportion of anti-inflammatory macrophages did not change significantly (**Fig. 1E and F**). Thus, we successfully demonstrated that CVB3 mediates pro-inflammatory macrophage polarization in a cellular model. Moreover, to assess whether CVB3 has the same polarizing effect on cardiac macrophages, we further examined the type of CVB3-mediated macrophage polarization in a mouse model. After 7 days of intraperitoneally injecting BALB/c mice with 5×10^7^ PFU of CVB3, heart tissues from the experimental and control groups were collected for testing. The immunofluorescence test showed that the CVB3 group had a large number of macrophages in the heart tissue and high CD86 expression, with strong red fluorescence surrounding the nucleus, while the control group had lower CD86 protein expression. However, CD206 was expressed at lower levels in the CVB3 and control groups, showing a faint green fluorescence (**Fig. 1G**). Thus, CVB3 can also promote M1-like polarization in cardiac macrophages in the mouse model.

### CVB3 leads to macrophage polarization by initiating metabolic reprogramming of the urea cycle

The polarization of macrophages is influenced by multiple factors, and a growing body of research has highlighted the critical role of metabolic reprogramming in macrophage polarization. Recent studies have found that the downregulation of citrulline by ASS1 in the urea cycle is a key factor in pro-inflammatory macrophage activation^[19]^, but this finding has not been validated in viral infection models. We demonstrated that CVB3 activates pro-inflammatory macrophages, so whether there is regulation of metabolic reprogramming in CVB3-induced pro-inflammatory macrophage activation. The immunofluorescence results showed that the heart tissue of mice injected with CVB3 exhibited stronger red and green staining; that is, more CD68 and ASS1 were expressed. In addition, ASS1 overlapped with CD68 and fluoresced yellow (**Fig. 2A**). This experiment suggested that ASS1 is indeed upregulated in cardiac macrophages. We also detected the expression levels of ASS1 in the hearts of CVB3-infected mice at different time points via immunohistochemistry (IHC), which also confirmed that CVB3 infection in mice can cause upregulated ASS1 expression in cardiac cells in the early stages of infection (**Supplementary Fig. 3**).

**Figure 2.**
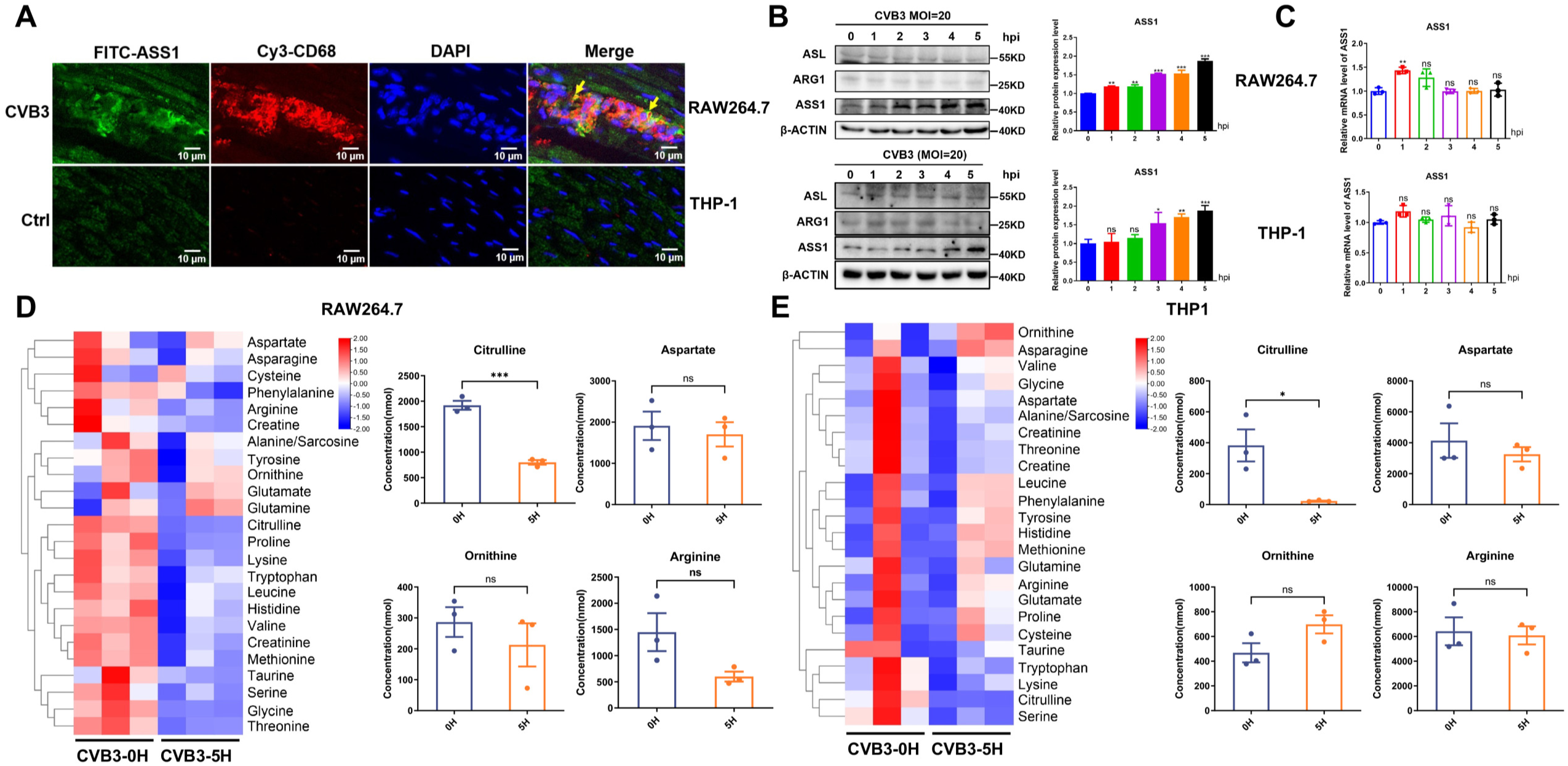
CVB3 leads to macrophage polarization by initiating metabolic reprogramming of the urea cycle. (A) After 7 days of intraperitoneal injection of BALB/c mice with 5×107 PFU of CVB3, immunofluorescence was used to detect ASS1 protein expression in mouse cardiac macrophages with VMC. The fluorescence intensity of cardiac macrophage marker protein Cy3-CD68 and urea cycle key enzyme FITC-ASS1 in the mouse VMC model was observed by fluorescence microscopy. FITC-labeled proteins appear green, Cy3-labeled proteins appear red, nuclei appear blue, and green light and red light overlap yellow. (B) Western blotting was used to detect the relative protein expression levels of ASS1, ARG1, and ASL in macrophages incubated with CVB3 (MOI = 20) for different durations. The intensities of ASS1 protein bands were quantified by ImageJ and data were presented as mean ± SD (*, P < 0.05; **, P < 0.01; ***, P < 0.001; ns, vs. the 0 hpi group). (C) Expression levels of ASS1 mRNA after CVB3 (MOI = 20) treatment of macrophages were measured by RT-qPCR. Values are shown as the mean ± SEM of three independent experiments (**, P < 0.01; ns, P > 0.05). (D and E) Hierarchical clustering (fragments per kilobase of transcript per million; k-means; Pearson’s correlation) of significantly changed genes at any indicated time point (n = 3). The absolute concentration of urea cycling-related amino acids in macrophages incubated with CVB3 (MOI = 20) for 5 h was determined by LC-MS compared with 0 h, data were presented as mean ± standard deviation (SD). (*, P < 0.05; ***, P < 0.001; ns, vs. the 0 hpi group).

To further validate our hypothesis, we evaluated the expression of enzymes of the urea cycle by western blotting and RT-qPCR analysis within 5 hours of CVB3 incubation. Expression levels of enzymes in CVB3-treated RAW264.7 and THP-1 cells were assessed. ASS1 protein expression was significantly increased, with arginase 1 (ARG1), OTC, and argininosuccinate lyase (ASL) remaining unchanged (**Fig. 2B**), while there was no obvious change in ASS1 mRNA, as well as in the mRNA levels of ARG1, OTC, and ASL (**Fig. 2C and Supplementary Fig. 4A and B**). To verify the changes in urea cycle-related amino acids under the action of CVB3, the metabolites of RAW264.7 and THP-1 cells incubated with CVB3 for 0 hours and 5 hours were subjected to liquid chromatography-mass spectrometry (LC-MS) amino acid-targeted detection and metabolomic analysis. The LC-MS assays demonstrated that citrulline levels were downregulated after 5 hours of virus incubation, while other relevant amino acids in the urea cycle remained unchanged (**Fig. 2D and E**). These results showed that CVB3 reprograms the urea cycle during M1-like activation and promotes citrulline depletion by upregulating ASS1 protein expression.

### Blocking citrulline depletion could rescue CVB3-induced pro-inflammatory macrophage polarization

Previously, we demonstrated that ASS1 plays a critical role as a key regulator of CVB3-induced metabolic reprogramming of the urea cycle at the onset of macrophage polarization, as well as metabolomics. Its upregulation at the protein level leads to low levels of citrulline in cells, which is beneficial for triggering macrophage polarization. We next investigated the role of citrulline metabolism in the immune response of CVB3-induced M1-like activation. CCK-8 assays confirmed that citrulline did not display cytotoxicity in macrophages (**Supplementary Fig. 5**). Citrulline addition dose-dependently inhibited the expression of M1 genes such as the marker CD86, the cytokines IL-6 and IL-1B, and pro-inflammatory genes in both RAW264.7 and THP-1 cells primed by CVB3 (**Fig. 3A, 3B and Supplementary Fig. 6**). We further confirmed these findings by flow cytometry; after CVB3 incubation, citrulline addition resulted in a reduced proportion of CD86^+^CD206^-^ cells, that is, fewer pro-inflammatory macrophages (**Fig. 3C and D**).

**Figure 3.**
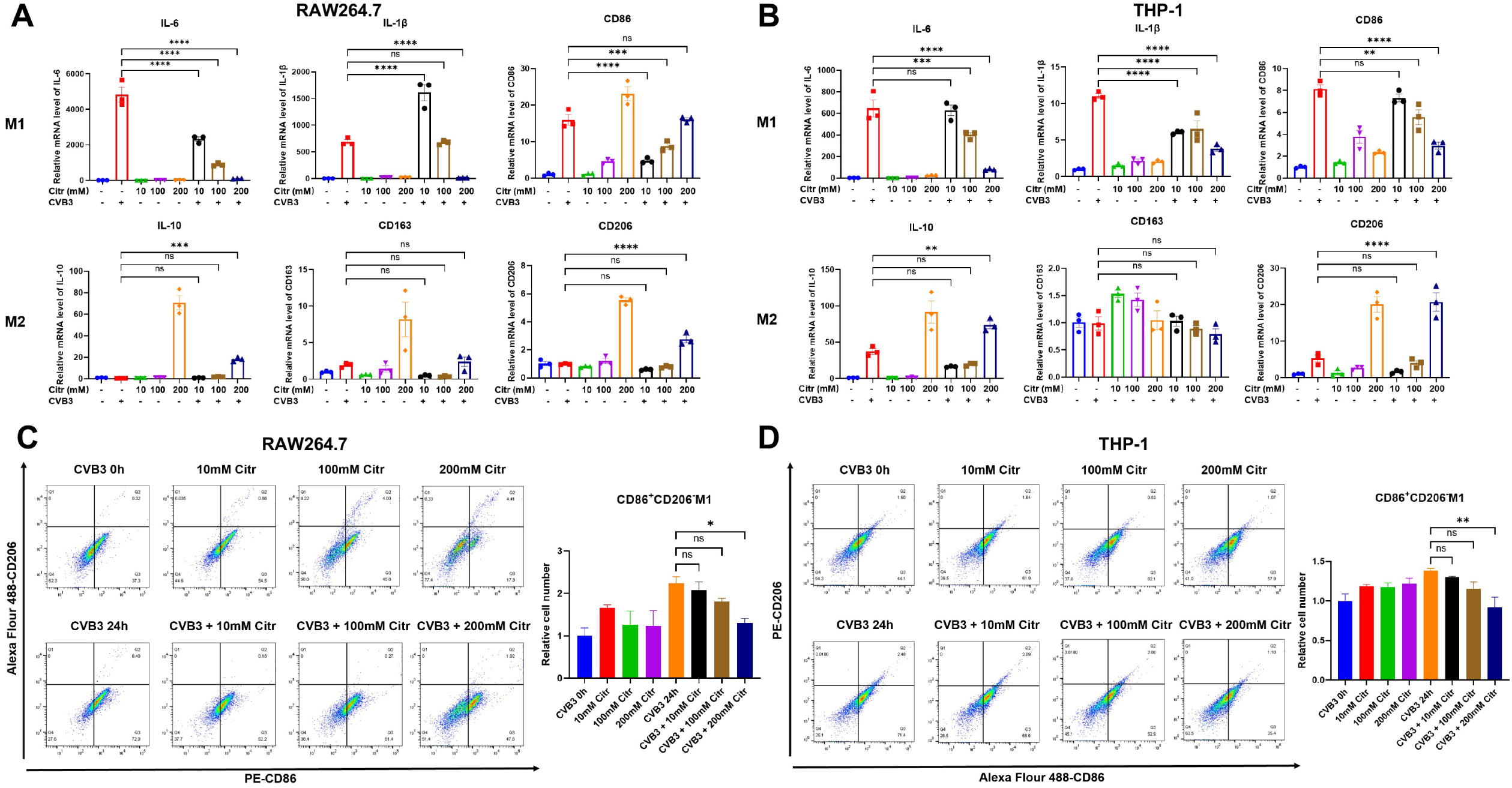
Exogenous citrulline supplementation could rescue CVB3-induced pro-inflammatory macrophage polarization. (A and B) Expression levels of M1/M2 macrophage marker mRNA after citrulline and CVB3 (MOI = 20) treatment of macrophages were measured by RT-qPCR. (C and D) Flow cytometry was used to detect the changes in CD86^+^CD206^-^M1 macrophages incubated with citrulline and CVB3 (MOI = 20). Values are shown as the mean ± SEM of three independent experiments. (*, P < 0.05; **, P < 0.01; ***, P < 0.001; ****, P < 0.0001; ns, P > 0.05).

### Citrulline could efficiently alleviate CVB3-induced myocarditis and was cardio-protective in CVB3-infected mice

To further investigate the role of citrulline metabolism in the immune response to CVB3-induced VMC in mammals, BALB/c mice were intraperitoneally injected with citrulline or Dulbecco’s modified Eagle’s medium (DMEM) every 2 days for 14 days, then with 5×10^7^ PFU of CVB3, during which period they were monitored for weight change and survival (**Fig. 4A**). Mice injected with citrulline survived and showed no obvious changes in body weight, indicating the low toxicity of citrulline. In addition, mice infected with CVB3 showed significant weight loss, and all CVB3-infected mice died by 7 days post-infection (dpi). By contrast, citrulline-treated CVB3-infected mice experienced less weight loss. Three-sevenths of CVB3-infected mice treated with citrulline were still alive at 7 dpi (**Fig. 4B and C**). H&E staining of heart tissues at 7 dpi, as well as IHC, proved that citrulline addition can alleviate the excessive inflammation and myocardial tissue damage in CVB3-infected mice and reduce ASS1 expression. Mice supplemented with citrulline appeared almost identical to the DMEM controls. Nevertheless, CVB3-infected mice showed significant myocardial injury and massive inflammatory cell infiltration in their hearts, while citrulline treatment significantly alleviated myocarditis and myocardial damage. Additionally, the average histopathological and immunohistochemical scores of the CVB3-infected group were significantly different from those of the citrulline-treated group (**Fig. 4D and E**). Moreover, the expression of pro-inflammatory macrophage markers and pro-inflammatory cytokines in the mouse hearts was significantly reduced (**Fig. 4F**). Citrulline potently reduced the viral capsid protein VP1 RNA and protein levels in CVB3-infected heart tissue (**Fig. 4G and H**). In summary, these data demonstrated that citrulline has a certain relieving and inhibitory effect on CVB3-induced myocarditis.

**Figure 4.**
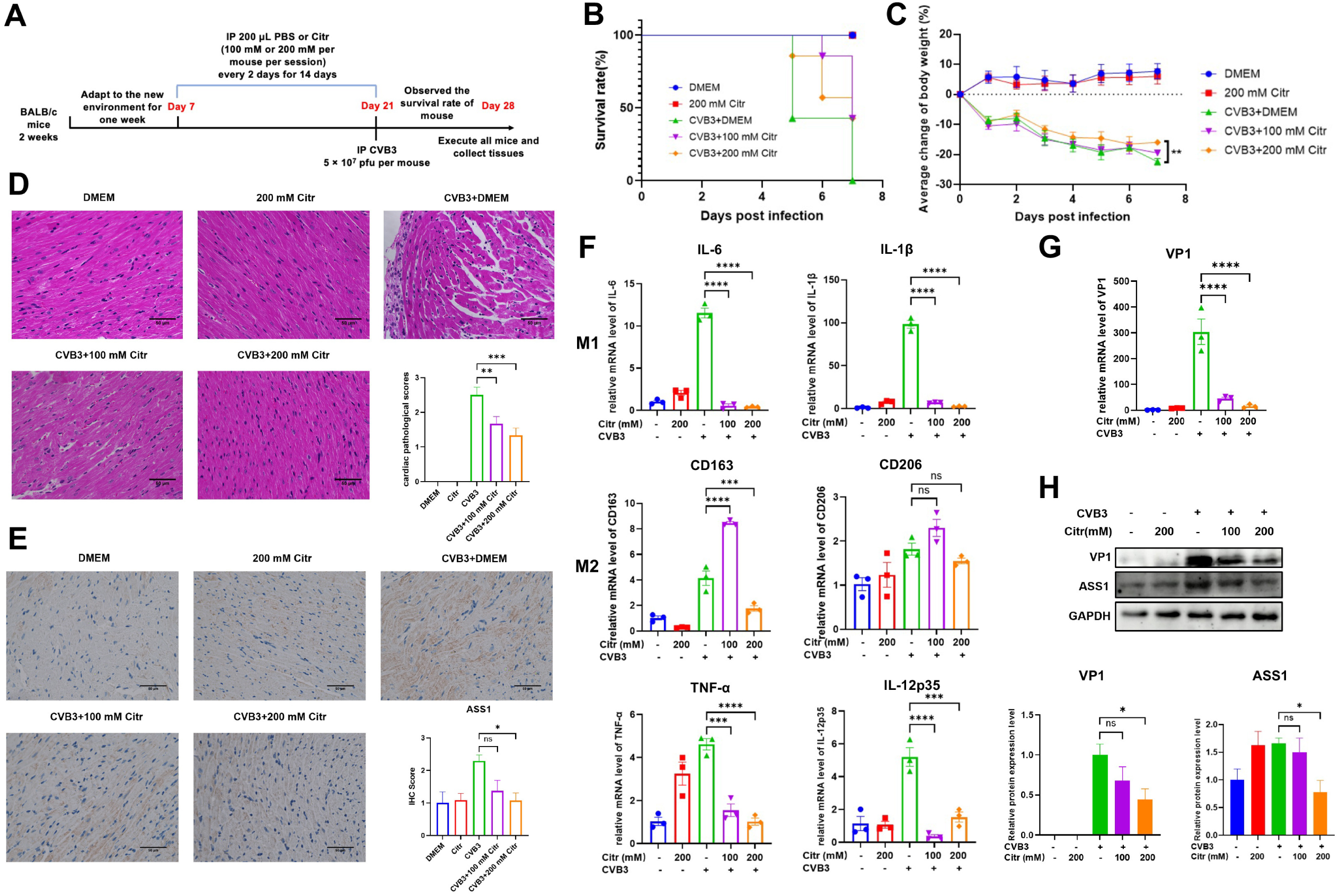
Citrulline could efficiently alleviate CVB3-induced myocarditis and was cardioprotective against CVB3-infected mice. (A) Schematic of the in vivo experiment. Mice were first adapted to the new environment, after which they were intraperitoneally injected with citrulline or DMEM every 2 days for 14 days, followed by intraperitoneal injection with 5×10^7^ PFU of CVB3 and humane execution after a further 7 days. (B and C) The survival rate and body weight data of the mice challenged with CVB3 (5×10^7^ PFU) were monitored daily until 7 dpi. (D) H&E staining and the pathology scores of heart sections from mice infected with CVB3 (5×10^7^ PFU) at 7 dpi. Scale bar = 50 μm. n = 6 in each group. (E) Representative IHC analysis of ASS1 in heart sections from mice infected with CVB3 (5×10^7^ PFU) at 7 dpi. Scale bar = 50 μm. n = 6 in each group. (F) Expression levels of pro-inflammatory cytokine and M1/M2 macrophage marker mRNA from three mice per group were measured by RT-qPCR. (G and H) Western blotting and RT-qPCR were used to detect the relative expression of viral capsid protein VP1 and ASS1 in CVB3-infected heart tissue. Statistical significance was based on one-way ANOVA (*, P < 0.05; **, P < 0.01; ***, P < 0.001; ****, P < 0.0001; ns, P > 0.05).

### CVB3 capsid protein VP2 interacts with ASS1 and inhibits its autophagic degradation

We found that ASS1 can be upregulated by CVB3. To further determine the regulatory relationship between CVB3 structural protein and ASS1, a coimmunoprecipitation experiment was performed, and the results showed that VP2 does interact with ASS1 (**Fig. 5A**). In addition, indirect immunofluorescence further demonstrated the interaction between VP2 and ASS1 and its colocalization in the cytoplasm (**Fig. 5B**). At the same time, we determined whether VP2 and other capsid proteins affect ASS1 expression. We selected HeLa cells, a CVB3 cell model, as the transfection vector due to the inherent difficulty in transfecting macrophages. As expected, increasing amounts of VP2 steadily increased the expression of ASS1 protein, while other capsid proteins did not affect ASS1 expression (**Fig. 5C**). In addition, citrulline treatment downregulated CVB3-induced ASS1 hyperexpression in RAW264.7 and THP-1 cells (**Fig. 5D**). Then, we hypothesized that the CVB3-induced metabolic reprogramming of macrophages may directly regulate ASS1. To understand the mechanism by which CVB3 upregulates ASS1, RAW264.7 cells were treated with CVB3 and the proteasome inhibitor MG132 and autophagy inhibitor 3-methyladenine (3-MA). The results showed that MG132 did not affect ASS1 expression, while 3-MA and CVB3 upregulated ASS1 and showed a synergistic effect (**Fig. 5E**), with the same results in VP2-treated HeLa cells (**Fig. 5F**). Huijun Zhang et al. found that foot-and-mouth disease virus’ structural protein VP3 interacts with histone deacetylase 8 (HDAC8) and promotes its autophagic degradation^[30]^. We speculated that the CVB3 capsid protein VP2 interacts with ASS1 and may inhibit its autophagic degradation. To verify our observation and hypothesis, western blotting was used to detect LC3B, and the results showed that the conversion of LC3-I to LC3-II was increased. CVB3 incubation and treatment with bafilomycin A1 (BafA1), an inhibitor of late-phase autophagy, also upregulated ASS1, Atg5, and LC3II, indicating that CVB3 may hijack autophagy (**Fig. 5E and F**). In summary, these findings suggest that CVB3 and its structural protein VP2 inhibit the degradation of autophagosomes and thus upregulate ASS1.

**Figure 5.**
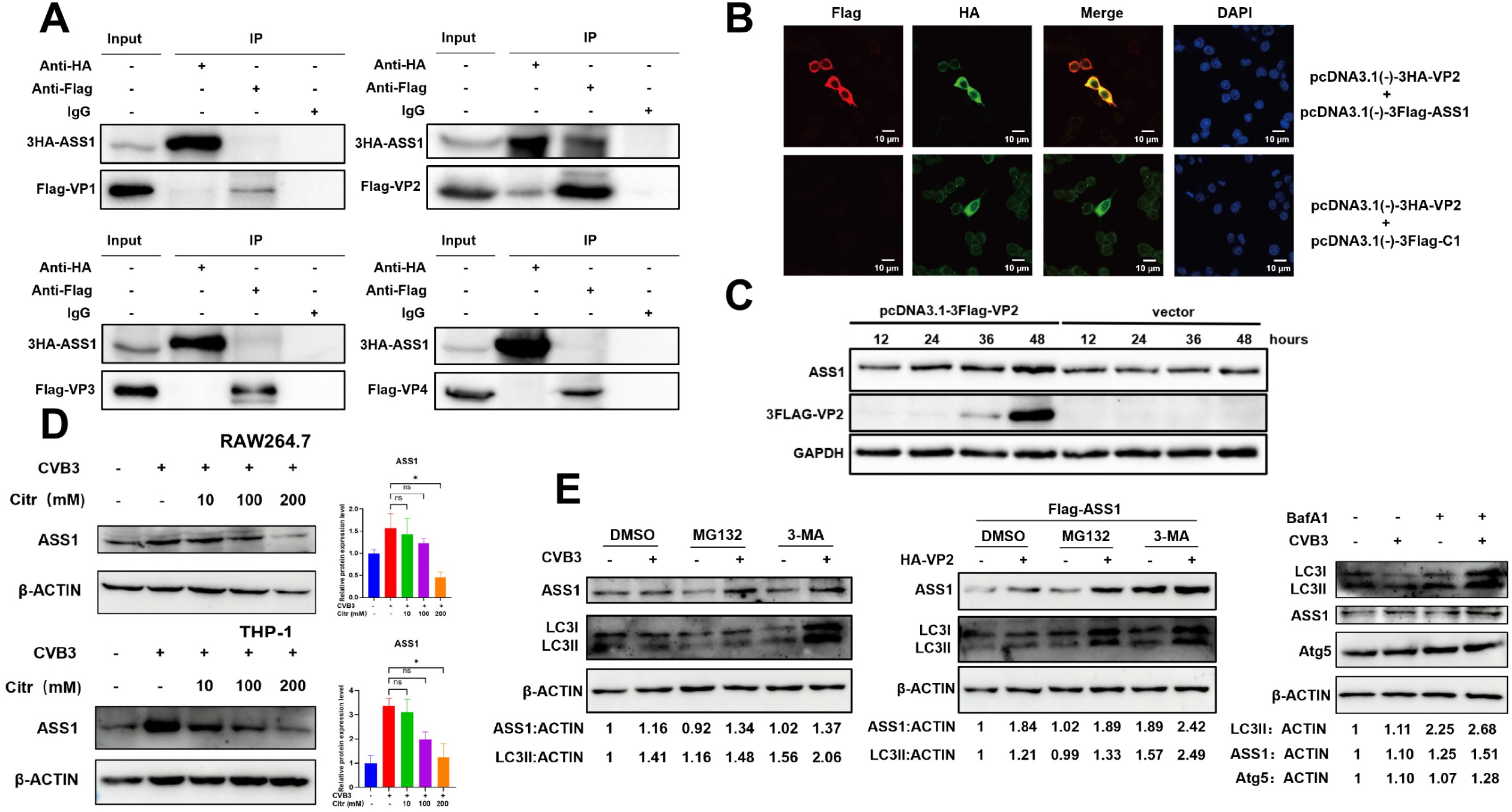
CVB3 capsid protein VP2 interacts with ASS1 and inhibits its autophagic degradation. (A) Co-IP verified the interaction between CVB3 capsid proteins VP1, VP2, VP3, and VP4 and ASS1 in 293T cells. (B) Indirect immunofluorescence staining was used to detect the colocalization between VP2 and ASS1 in HeLa cells by confocal microscopy. (C) The effect of CVB3 capsid protein on the expression of ASS1 protein was detected by western blotting. (D) Western blotting was used to determine ASS1 protein expression in macrophages incubated with CVB3 (MOI = 20) in the absence (-) or presence of increasing amounts of citrulline. (E) Western blotting was used to detect ASS1 and LC3B in RAW264.7 cells treated with DMSO, 10 μM MG132, and 5 mM 3-MA for 8 h or 100 nM BafA1 for 12 h, with CVB3 (MOI = 20) treatment for 5 h. (F) HeLa cells were transfected with Flag-tagged ASS1 plasmid with or without HA-tagged VP2 plasmid; 40 h later, the cells were treated with DMSO, 10 μM MG132, and 5 mM 3-MA for 8 h or 100 nM BafA1 for 12 h. Statistical significance was based on one-way ANOVA (*, P < 0.05; **, P < 0.01; ***, P < 0.001; ns, P > 0.05).

## Materials and methods

### Ethics statement

The CVB3 virus strain was stored in a biosafety level 2 laboratory, and all experiments met the requirements of the experimental guidelines of Nanchang University. Animal experiments were performed in accordance with the guidelines of the Laboratory Animal Ethics Committee of Nanchang University (Approval ID NCU-2021-0010).

### Cells and viruses

RAW264.7 cells, HeLa cells, and human embryonic kidney (HEK293T) cells were maintained in DMEM (Solarbio, China) supplemented with 10% fetal bovine serum (Gibco, USA). THP-1 cells were maintained in RPMI-1640 (Solarbio, China) supplemented with 10% fetal bovine serum (Gibco, USA), Thereafter, the cells were differentiated into non-polarized (M0) macrophages using 75 ng/mL phor-bol 12-myristate 13-acetate (PMA, GlpBio, USA) for 24 h, washed, and re-incubated for 24 h before use in experiments. All the cells were cultured in a 5% CO_2_ incubator at 37 °C. CVB3 (Nancy strain; GI:323432) was propagated and amplified in HeLa cells and stored at −80 °C for further usage.

### Antibodies and reagents

PE anti-mouse CD86 antibody, Alexa Fluor® 488 anti-mouse CD206 antibody, Alexa Fluor® 488 anti-human CD86 antibody, PE anti-human CD206 antibody, and PE anti-mouse/human CD11b antibody were purchased from BioLegend (USA); anti-CD86 (rabbit), anti-CD206 (rabbit), anti-CD68 (rabbit), CY3-tyramide (tsa), and FITC-tyramide (tsa) were purchased from Servicebio (China); anti-ASS1 (rabbit) antibody was purchased from Abcam (USA); anti-ASL (mouse) and anti-ARG1 (rabbit) antibodies were purchased from PTM BIO (China); anti-enterovirus VP1 antibody was purchased from Dako (Denmark); anti-β-actin (mouse), anti-LC3B (rabbit), anti-Atg5 (rabbit), and TRITC-goat anti-mouse IgG antibodies were purchased from Proteintech (China); anti-FLAG (rabbit) antibody was purchased from Nalgene (USA); HA-tag rabbit mAb was purchased from Cell Signaling Technology (USA); FLAG-tag mouse mAb was purchased from Sigma (USA); goat anti-rabbit IgG H&L/HRP was purchased from Bioss (China); protein A/G PLUS-agarose was purchased from Santa Cruz Biotechnology (USA); Alexa Fluor 488-goat anti-rabbit IgG was purchased from Abmart (China); HRP-conjugated AffiniPure goat anti-rabbit IgG (H+L) and HRP-conjugated AffiniPure mouse anti-rabbit IgG (H+L) were purchased from Boster (USA). PMA and the CCK-8 assay were purchased from GlpBio (USA); TRIzol, PrimeScript^TM^RT Master Mix and TB Green® Premix Ex Taq™ were purchased from Takara (Japan); Triton-X100 and L-citrulline were purchased from Solarbio (China); MG132 was purchased from Santa Cruz Biotechnology (USA); and 3-MA and BafA1 were purchased from MCE (USA).

### Plasmid construction, site-directed mutagenesis, and transfection

The ASS1 (Gene ID: 445) genes were amplified by PCR from the cDNA of HeLa cells and then cloned into the pcDNA3.1(-)-3HA vector and pcDNA3.1(-)-3Flag vector, generating the expression plasmids pcDNA3.1(-)-3HA-ASS1 and pcDNA3.1(-)-3Flag-ASS1. cDNA fragments encoding all the viral proteins derived from CVB3 were amplified by PCR and then cloned separately into the vectors pCAG-FLAG and pcDNA3.1(-)-3Flag. All the primers are listed in Table S1. All plasmids were transfected into cells using PEI 40K transfection reagent (Servicebio, China) according to the manufacturer’s instructions.

### Cell viability assay

Cell viability was analyzed using a CCK-8 assay (GlpBio, USA). A total of 7000 cells per well were seeded in a 96-well plate and, after nearly 12 h of incubation, incubated with citrulline for 12 h. Then, 100 μL of CCK-8 solution was added per well and incubated for 1.5 h in a 5% CO_2_ incubator at 37 °C. A microplate reader was then used to measure OD at 450 nm.

### RNA isolation and RT-qPCR

Total RNA was extracted using TRIzol (Takara), and 500 ng of RNA was reversed to cDNA with PrimeScript^TM^RT Master Mix (Takara). RT-qPCR was performed with 100 ng of the above cDNA as a template, with at least three replicates. Relative mRNA expression levels were calculated using actin as an internal control and normalized to those of the control groups.

### Flow cytometry

Cells were fixed with 4% paraformaldehyde (Servicebio) for 15 min, then treated with PBS containing 0.1% Triton-X100 (Solarbio) for 5 min (mouse CD206 is a transmembrane antigen), washed, and incubated with relevant antibodies for 30 min on ice, then washed again and analyzed on the flow cytometer (BD FACSVerse). FlowJo v10.6.2 software (Tree Star) was used to analyze the data.

### Immunofluorescence assay

After the heart tissue was sectioned, antigen retrieval was performed using ethylenediaminetetraacetic acid (EDTA; pH 8.0; Servicebio), and then the sections were placed in a 3% hydrogen peroxide solution to block the endogenous peroxidase. After washing with PBS, 3% BSA (Solarbio) was added for blocking for 30 min, then the blocking solution was removed, the first primary antibody was added, and the sample was incubated overnight at 4 °C. Subsequently, the sample was washed, the corresponding HRP-conjugated secondary antibody was added, and incubation took place at room temperature in the dark for 1 h. Next, the corresponding tsa dye was added, and then antigen retrieval was performed again; the second group of primary antibodies, secondary antibodies, and the corresponding tsa dye were added, and finally, the nuclei were counterstained with DAPI and observed with a FLUOVIEW FV1000 confocal laser scanning microscope (Olympus).

### H&E staining and IHC

Mouse heart tissues were isolated and fixed in 10% formaldehyde for 24 h at room temperature. Fixed tissues were dehydrated and paraffin-embedded. For H&E staining, slices were rehydrated and stained with hematoxylin and eosin according to the manufacturer’s instructions. Myocarditis pathology score was assessed according to the following criteria: 1 = presence of inflammatory cell foci between myocytes or around individual myocytes; 2 = 5-10% myocardial section involvement; 3 = 10-20% myocardial section involvement.

For IHC, fixed tissues were deparaffinized and rehydrated, followed by antigen retrieval, washed three times with PBS (pH 7.4), then treated with anti-ASS1 antibody (Abcam), incubated with secondary antibody, stained with 3,3-diaminobenzidine, back-stained with hematoxylin, and viewed with a microscope (Invitrogen, EVOSM5000) after mounting. Image J and the IHC Profiler plug-in were used for immunohistochemical scoring, and finally, the data were normalized.

### Co-immunoprecipitation (co-IP) and western blotting

For western blotting, cells were lysed in lysis buffer (150 mM NaCl, 20 mM Tris HCl, 0.1% NP-40, pH 7.4) and then heated for 10 min at 100 °C. Equal amounts of protein were separated by an sodium dodecyl sulfate polyacrylamide gel electrophoresis (SDS-PAGE) gel and transferred to an Nitrocellulose (NC) membrane, then the membranes were blocked with 5% skim milk for 2 h at room temperature, incubated overnight at 4 °C with the primary antibody, and then incubated with the secondary antibody for 2 h at room temperature. Finally, the protein was visualized with Super ECL Detection Reagent (YEASEN).

For Co-IP, HEK293T cells were transfected with the indicated plasmids for 48 h; then, cells were lysed with the Native Lysis Buffer containing 1% Phenylmethylsulfonyl fluoride (PMSF, Solarbio) and the supernatant of the lysate was collected after centrifugation and divided into four aliquots. One aliquot was used as the input sample, and the remaining three aliquots were immunoprecipitated overnight at 4 °C with the indicated antibody mixed with Protein A/G PLUS-Agarose (Santa Cruz Biotechnology). Then, the beads containing the immunoprecipitation complex were washed three times with the Native Lysis Buffer containing 1% PMSF. Both input samples and immunoprecipitation complexes were lysed with lysate (150 mM NaCl, 20 mM Tris HCl, 0.1% NP-40, pH 7.4) and then heated for 10 min at 100 °C, followed by western blotting.

### Indirect immunofluorescence microscopy

After 48 h of VP2 transfection with ASS1 or vector plasmid, the anti-Flag and anti-HA primary antibodies were incubated overnight at 4 °C. Then, cells were washed three times at room temperature with PBST. The fluorochrome-conjugated secondary antibodies (Alexa Fluor 488-goat anti-rabbit IgG, 1:200; TRITC-goat anti-mouse IgG, 1:200) were incubated with cells for 2 h in the dark, extensively washed with PBST, and counterstained with DAPI. The stained slides were visualized using a FLUOVIEW FV1000 confocal laser scanning microscope (Olympus) after fixation.

### LC-MS analysis of amino acid metabolism

RAW264.7 and THP-1 cells were incubated with CVB3 for 0 h or 5 h. After washing, the cells were resuspended with 1 mL of pre-chilled methanol: acetonitrile: water = 2:2:1 and stored in a −80 °C freezer. LC-MS metabolomics detection was performed by Applied Protein Technology Co., Ltd.

### Citrulline supplementation

Citrulline was dissolved in DMEM or RMPI 1640 medium and filtered with a 0.22 μm filter (BIOFIL). For cell experiments, cells were incubated with citrulline for 12 h before being incubated with the virus. For animal experiments, mice were intraperitoneally injected with citrulline every 2 days for 2 weeks prior to CVB3 infection.

### Statistical analysis

Data were analyzed for statistical significance by Student’s t test or one-way analysis of variance (ANOVA). Statistics were performed using GraphPad Prism 9.5.1 and ImageJ 1.52a. Graphs display mean values, with error bars representing the standard error of the mean (SEM) in all graphs. Differences were considered statistically significant at P < 0.05 (*, P < 0.05; **, P < 0.01; ***, P < 0.001; ****, P < 0.0001; ns, P > 0.05).

## DISCUSSION

In this work, we preliminarily clarified the initiation mechanism of CVB3-induced VMC in cell models and animal models. Here, CVB3 capsid protein VP2 inhibited cellular autophagy, resulting in urea cycle metabolic reprogramming, manifested by the eventual upregulation of ASS1 and citrulline depletion, which, in turn, activates macrophage pro-inflammatory polarization, resulting in the occurrence of acute myocarditis (**Fig. 6**). In addition, we preliminarily evaluated the therapeutic potential of citrulline for VMC and found that citrulline supplementation not only rescues CVB3-induced macrophage polarization but also effectively alleviates the disease manifestations and pathogenicity of CVB3-induced VMC. These results provide a new research direction for the occurrence and development mechanisms of VMC, suggesting that ASS1 may be a potential target for the treatment of this disease.

**Figure 6.**
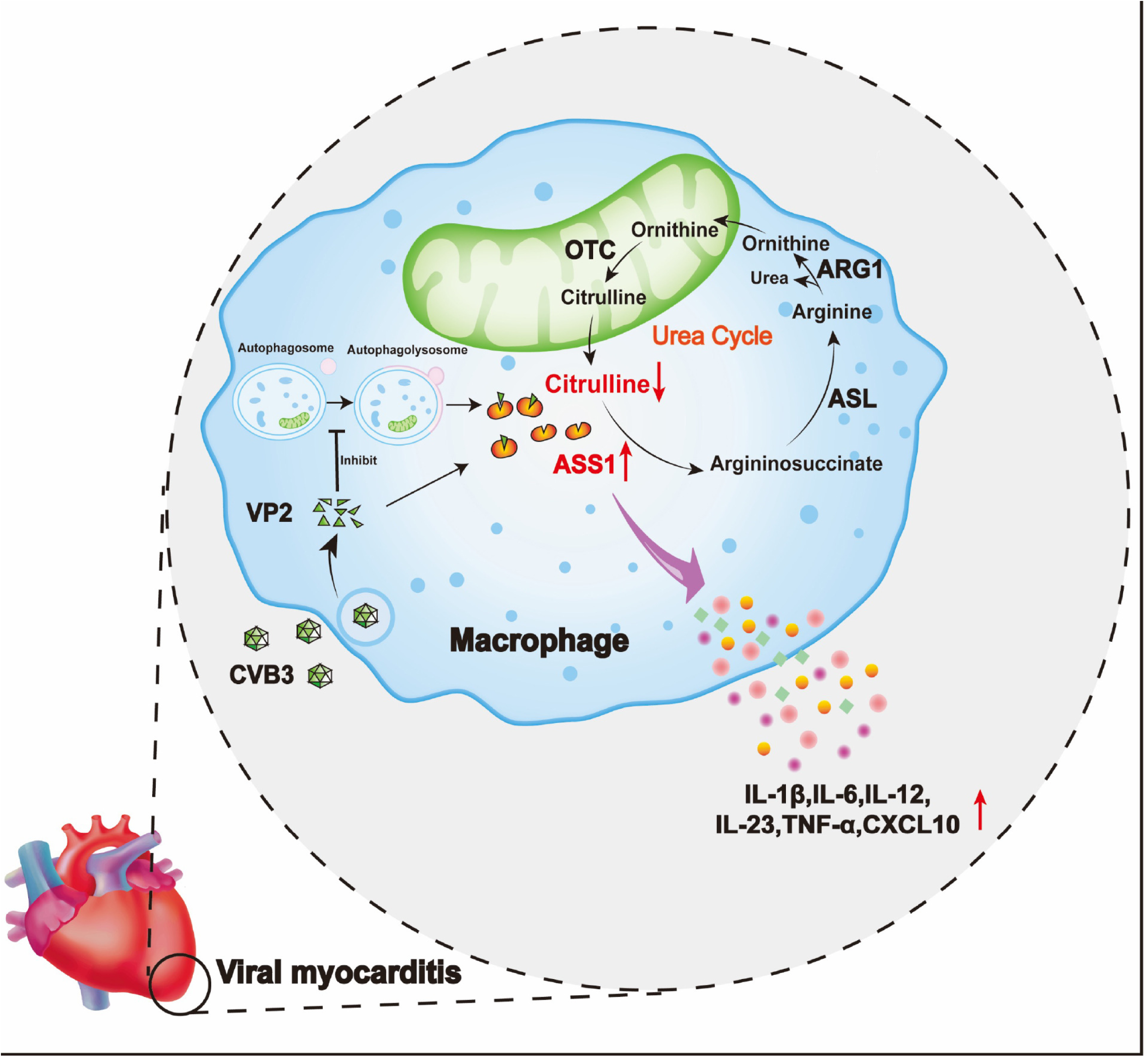
Schematic diagram of CVB3 activating macrophage pro-inflammatory polarization and thereby promoting VMC. CVB3 capsid protein VP2 inhibits ASS1 autophagic degradation and upregulates ASS1, triggers urea cycle metabolic reprogramming, activates macrophage pro-inflammatory polarization, and promotes the development of VMC.

CVB3 is an enterovirus that causes acute myocarditis. VP1, VP2, VP3, VP4 are the four capsid proteins encoded by CVB3. By binding to the CAR/CD55/DAF receptor on the host cell membrane, the VP4 and VP1 proteins hidden within the nucleocapsid are exposed, thereby promoting the encapsulation and uptake of CVB3 by the host cell^[31]^. Currently, the relationship between the CVB3 capsid protein and the host is mainly studied via VP1, which is considered a key capsid protein that can interfere with the host cell cycle and mediate apoptosis^[32–34]^. In addition, the relationship between capsid proteins VP2, VP3, and VP4 and their hosts has attracted increasing attention. VP2 is thought to be a key pathogenic factor involved in VMC, primarily involved in mediating apoptosis in host cells^[25, 35, 36]^. When VP3 is mutated, polyamine production is reduced and can affect viral attachment^[37]^, and mutations may affect virulence in mouse models^[38]^. VP4 is thought to be the target site for CVB3 to stimulate the release of antibodies from the interferon-alpha (IFN-α) signaling pathway^[39]^. Furthermore, VP1 and VP2 are considered to be the key proteins that promote macrophage activation^[40]^, and mutations of VP2 and VP3 can reduce cardiovirulence and cause CVB3 immune evasion^[41]^. These results suggested that the capsid protein of CVB3 may be an important causative factor of inflammation. Similarly, we found that VP2 may also be a key pathogenic factor involved in VMC, and its upregulation of ASS1 causes urea cycle metabolic reprogramming to promote VMC development.

Metabolic reprogramming of the urea cycle leads to tumor progression. For example, ASS1 is downregulated in hepatocellular carcinoma and pancreatic cancer^[42, 43]^ but upregulated in ovarian, colorectal, and gastric cancers^[44–46]^ and can produce arginine, making ASS1 a potentially effective therapeutic target for these cancers. Previous studies have shown that ASS1 is a pro-inflammatory gene^[47]^; likewise, we found that in the early stage of CVB3 infection, ASS1 was elevated and depleted its substrate citrulline, thereby activating pro-inflammatory macrophages, while the exogenous supplementation of citrulline inhibited the expression of ASS1, thereby limiting the activation of macrophages and the release of pro-inflammatory cytokines and alleviating myocardial injury caused by CVB3. To demonstrate the interconnection between CVB3 and the host, and that ASS1 was upregulated at the protein level, while transcription was not affected. We hypothesized that there may be an interaction between CVB3 capsid proteins involved in inflammation and ASS1. As we suspected, there was an interaction between ASS1 and VP2. CVB3 upregulates ASS1, and VP2 also upregulates ASS1, which also suggests that ASS1 may be a potential target for VMC. However, due to the difficulty in transfecting macrophages, this experiment was validated in a CVB3 cell model: HeLa cells.

In addition, we were pleasantly surprised to discover the potential of citrulline in the treatment of VMC. We found that citrulline can downregulate ASS1, affect urea cycle metabolism, inhibit macrophage activation, and rescue CVB3-induced macrophage pro-inflammatory polarization, thereby effectively alleviating the disease manifestations of VMC. Previous studies have also demonstrated the potential of citrulline in treating cardiovascular and inflammatory diseases^[48, 49]^. However, we did not get the best results, and we speculated that long-term high-dose citrulline intake may hinder its metabolism in mice^[19]^. We thus plan to use low doses of citrulline as an adjunct to antiviral drugs in treating VMC in the hope of obtaining better therapeutic effects.

Previous studies have shown that CVB3 infection promotes the formation of autophagosomes but disrupts the fusion mechanism required for normal autophagic flux, inhibiting autophagosome binding to lysosomes, thereby hijacking cellular autophagy to promote viral replication^[50]^. In addition, Kun Liu et al. found that macrophage autophagy defects may promote pro-inflammatory macrophage polarization, which may be the basis of inflammatory diseases^[51]^. Huijun Zhang et al. found that the foot-and-mouth disease virus structural protein VP3 interacts with HDAC8 and promotes its autophagic degradation^[30]^. In this work, we found that CVB3 and its capsid protein VP2 hijacked cell autophagy, upregulated ASS1, and depleted citrulline, thereby promoting the pro-inflammatory polarization of macrophages and the occurrence and development of VMC. However, the inhibition of macrophage autophagy by CVB3 needs to be further validated by other methods, such as indirect immunofluorescence and electron microscopy, and future studies are warranted to determine whether the knockdown of ASS1 inhibits CVB3-induced pro-inflammatory macrophage activation and whether CVB3 attenuates the pathogenicity of VMC in mice with ASS1 knockout.

There is currently no specific treatment for VMC caused by CVB3. In recent years, the development of treatment regimens for macrophage polarization and metabolic reprogramming has become a new focus. Pengcheng Yan et al. found that dapagliflozin, as a sodium-glucose co-transporter 2 (SGLT-2) inhibitor, increased anti-inflammatory macrophage polarization after VMC, thereby reducing cardiac damage^[52]^. Recent studies and the present study indicate that ASS1 is expected to be used as a breakthrough in the treatment of VMC caused by CVB3 and that ASS1 inhibitors targeting VP2 are to be found to promote ASS1 degradation, inhibit pro-inflammatory macrophage polarization, and reduce myocardial inflammatory infiltration. Our experiments have also confirmed that citrulline may also have therapeutic potential from the perspective of interfering with amino acid metabolism.

## Compliance with Ethical Standards

### Funding

This study was supported by the National Natural Science Foundation of China (82203032, 32260193 and 32060040), Natural Science Foundation of Jiangxi Province (20202BAB206062), Training Plan for Academic and Technical Leaders of Major Disciplines in Jiangxi Province-Youth Talent Project (20212BCJ23036), Project for high and talent of Science and Technology Innovation in Jiangxi “Double-Thousand Talents Program of Jiangxi Province” (jxsq2023301110 and jxsq2023201019) and the National Innovation and Entrepreneurship Training Program for college students (202110403093 and 202310403042).

## Acknowledgement

The authors would like to express their gratitude to EasytoEdit (https://www.yijisci.com/) for the expert linguistic services provided.

## Ethical approval

Not applicable.

## Data availability

The datasets generated during and analysed during the current study are available from the corresponding author on reasonable request.

## Code availability

Not applicable.

## Consent to participate

Not applicable.

## Consent for publication

Not applicable.

## Author contributions

Qiong Liu and Xiaotian Huang conceived and designed the experiments; Qiong Liu, Yinpan Shang, Ziwei Tao, Xuan Li, Hanchi Zhang, and Lu Shen performed the experiments. Hanchi Zhang, Ziwei Tao, Zhili Liu, Zhirong Rao, Xiaomin Yu, Yanli Cao, and Lingbing Zeng analyzed the data; Qiong Liu, Yinpan Shang, Xuan Li, and Xiaotian Huang wrote the manuscript.

---

**Supplementary Figure 1.** H&E staining of heart sections of mice infected with 5×10^7^ PFU of CVB3 at several hours post-infection. Scale bar = 50 μm.

**Supplementary Figure 2.** The expression level of VP1 mRNA in CVB3-infected and CVB3-incubated RAW264.7 cells was detected by RT-qPCR (MOI = 20).

**Supplementary Figure 3.** Representative IHC analysis of ASS1 in heart sections from mice infected with CVB3 (5×10^7^ PFU) at several hours post-infection. Scale bar = 50 μm. n = 6 in each group. Statistical significance was based on one-way ANOVA (*, P < 0.05; ****, P < 0.0001; ns, P > 0.05).

**Supplementary Figure 4.** (A and B) RT-qPCR was used to detect the relative mRNA expression levels of ARG1, ASL, ORNT1, and OTC in macrophages incubated with CVB3 (MOI = 20) for different durations. Statistical significance was based on one-way ANOVA (*, P < 0.05; **, P < 0.01; ***, P < 0.001; ns, P > 0.05).

**Supplementary Figure 5.** (A and B) The CCK-8 assay was used to examine the effect of citrulline on the activity of RAW264.7 and THP-1 cells.

**Supplementary Figure 6.** (A and B) Expression levels of pro-inflammatory genes and chemokine mRNA after citrulline and CVB3 (MOI = 20) treatment of macrophages were measured by RT-qPCR. Statistical significance was based on one-way ANOVA (*, P < 0.05; ****, P < 0.0001; ns, P > 0.05).

